# How Good is AlphaFold3 at Ranking Drug Binding Affinities?

**DOI:** 10.1101/2025.05.27.656341

**Authors:** Xin Hong, Bowen Gao, Yinjun Jia, Wenyu Zhu, Qixuan Chen, Xiaohe Tian, Zhenyi Zhong, Jianhui Wang, Yanyan Lan

## Abstract

Accurate affinity ranking of small molecules is pivotal for drug discovery. We investigate whether structure prediction models like AlphaFold3, pretrained on protein-ligand interactions, can address this task. Zero-shot evaluation of Protenix (an AlphaFold3-like model) demonstrates superior prioritization of active compounds over conventional scoring functions and state-of-the-art deep learning models. By further fine-tuning Protenix on structure-agnostic protein-ligand bioactivity data from ChEMBL and BindingDB, we develop AlphaRank that predicts pairwise affinity relationships. AlphaRank achieves prediction accuracy comparable to computationally intensive free energy perturbation (FEP+) workflows on standard benchmarks, while requiring substantially less computational resources. Our findings highlight the emergent potential of AlphaFold3-derived models in affinity ranking tasks and emphasize the necessity for targeted methodological exploration to fully harness their capabilities in drug discovery applications.

## 1. Introduction

Accurate small molecule affinity ranking is fundamental to drug discovery, particularly in virtual screening and lead optimization. While free energy perturbation (FEP+) remains the most reliable method, its prohibitive computational cost limits practical use. Recent breakthroughs in protein structure prediction, exemplified by AlphaFold3 (Abramson et al., 2024), have achieved unprecedented accuracy in modeling protein-ligand interaction patterns. This prompts a pivotal question: can such interaction-aware models transcend structural prediction to address the critical challenge of efficient affinity ranking?

We first propose a zero-shot strategy for affinity ranking by co-folding target protein sequences with pairs of small molecule SMILES using Protenix (Team et al., 2025), an AlphaFold3-like model. Remarkably, this approach reveals a competitive binding mechanism, that higher-affinity molecules preferentially occupy the real binding pockets. Evaluated on the FEP benchmark, this method surpasses traditional energy-based scoring functions like MM-GBSA and rivals task-specific deep learning models. These results demonstrate that structural pretraining in AlphaFold3-derived models inherently captures essential protein-ligand interaction patterns, enabling competitive affinity discrimination without explicit affinity-focused optimization.

To enhance alignment between structural features and affinity ranking, we develop AlphaRank by integrating Protenix’s PairFormer outputs with a lightweight network trained on approximately 30,000 structure-agnostic bioactivity entries from ChEMBL and BindingDB. Two input strategies are explored: **AlphaRank**_**triplet**_ predicts relative affinity by jointly processing protein–ligand–ligand triplets, while **AlphaRank**_**pair**_ independently evaluates individual protein–ligand pairs through separate co-folding. Finally, combining both strategies, **AlphaRank**_**ensemble**_ achieves state-of-the-art accuracy on both test sets, surpassing all learning-based baselines and closely approaching the performance of the physics-based gold standard FEP+. Crucially, this approach eliminates structural prerequisites by relying solely on sequence inputs, offering broad applicability in structure-agnostic drug discovery scenarios.

Analysis reveals that affinity ranking accuracy varies by task difficulty, i.e. pairs with large affinity gaps are reliably distinguished, while subtle differences remain challenging. Protein-level performance strongly correlates with structural confidence scores (pLDDT), confirming that prediction quality drives ranking capability. This suggests dual optimization paths for AlphaRank: refining discrimination of marginal affinity differences through adversarial training, and integrating known structural anchors when pLDDT indicates low prediction reliability.

## 2. Related Works

Traditional physics-based affinity prediction methods like FEP+ (Wang et al., 2015) offer high accuracy prediction through alchemical transformation-based free energy calculations but demands intensive molecular dynamics sampling, making it impractical for large-scale use. MM-GB/SA (Genheden & Ryde, 2015) is an alternative faster method but is less precise. Recent deep learning approaches have shown promise in affinity ranking with distinct strategies: PBC-Net (Yu et al., 2023) leverages ligand pair feature differences via graph networks, EHIGN (Yang et al., 2024) encodes multi-type protein-ligand interactions, and LigUnity (Feng et al., 2025) combines screening and listwise ranking data for improved predictions. Recent attempts explore AlphaFold3’s confidence metrics for affinity ranking, such as using confidence score ipTM as affinity proxies (Shamir & London, 2025). However, the systematic adaptation of AF3-derived models for accurate, generalizable affinity prediction remains an open question.

## 3. Method

### 3.1. Zero-shot Competitive Binding

While the previous zero-shot method (Shamir & London, 2025) leveraging interface predicted TM-scores (ipTMs) demonstrate baseline utility in affinity ranking, these structural confidence metrics lack explicit design for affinity ranking. We address this mismatch by developing a competitive binding strategy that incorporates affinity ranking’s essential inductive biases.

Specifically, for each target protein *p*, we rank a ligand pair (*l*_1_ and *l*_2_) by inputting their SMILES strings alongside the protein sequence and preprocessed MSA into Protenix. The model jointly folds the triplet (*p, l*_1_, *l*_2_) into a complex to simulate competitive binding. To assess ranking accuracy, we first define a reference pocket *P*_ref_ using residues within 6Å of the ligand in the co-crystal structure. Predicted binding pockets for each ligand are then extracted in the similar way, and then aligned from the predicted structure to the reference via sequence mapping. The affinity ranking accuracy is determined by whether the higher-affinity ligand’s pocket achieves greater spatial overlap (IoU) with the reference, where IoU is defined as:

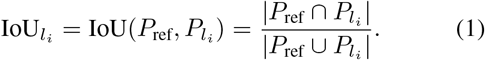

A higher 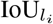 indicates greater overlap with the known binding site and is used as a proxy for binding quality.

To account for structure prediction uncertainty, we fold *K* times per (*p, l*_1_, *l*_2_) triplet, and ensemble the results through an averaged normalized score:

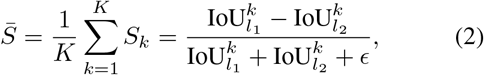

where *ϵ* is a small constant to prevent division by zero, and 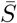 ranges from − 1 to 1, with positive values favoring *l*_1_ and negative values favoring *l*_2_.

### 3.2. Fine-tuning with PairFormer Features

Our next target is to improve affinity ranking by fine-tuning AF3-like models on protein-ligand bioactivity data. We focus on intermediate features from the PairFormer module, the core interaction engine which produces single **s** and pairwise (**z**) representations. A lightweight prediction head processes these features through two input paradigms: triplet-based co-folding AlphaRank_triplet_ for direct ligand competition analysis and AlphaRank_pair_ for pair-based independent scoring.

#### 3.2.1. Triplet-based Fine-tuning

Following the zero-shot setting, AlphaRank_triplet_ encodes the triplet (*p, l*_1_, *l*_2_) and then slices ligand atom features 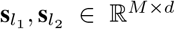 and interaction features between protein-ligand 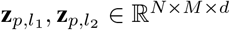 from PairFormer outputs’ single feature **s** and pair feature **z**. These features undergo average pooling to produce condensed vectors 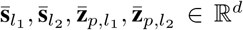 which are concatenated and processed by an MLP followed with a sigmoid function *σ*:

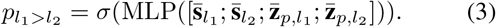

The model is trained with a binary cross-entropy loss:

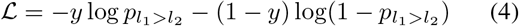

where *y* = 1 if *l*_1_ is the stronger binder and 0 otherwise. This training objective enables direct learning of comparative binding behavior from labeled bioactivity data to improve the ability of affinity ranking.

#### 3.2.2. Pairwise-based fine-tuning

AlphaRank_pair_ processes protein-ligand pairs (*p, l*_1_) and (*p, l*_2_) independently through the PairFormer module, generating ligand atom embeddings 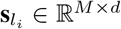 and pairwise interaction features 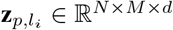. Averaging pooling condenses these into fixed-dimensional vectors 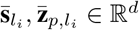, concatenated and transformed by an MLP:

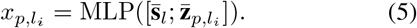

To enable the model to learn to rank affinities, training employs a pairwise ranking loss from RankNet (Burges et al., 2005):

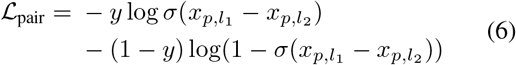

where *y* = 1 if *l*_1_ is the stronger binding molecules.

### 3.3. Training data curation

#### 3.3.1. Data collection

We curated activity and binding data for single-protein targets from ChEMBL35 (Mendez et al., 2018) and BindingDB (Gilson et al., 2016), processing each dataset separately. Compounds were desalted, and only those with molecular weights between 100 and 800 Da were retained, provided they contained no uncommon elements or long unbranched chains exceeding six atoms. Activity measurements such as IC_50_, EC_50_, K_*i*_, and K_*d*_ were standardized by conversion to negative logarithmic molar values. We processed the data to eliminate redundancy by deduplicating entries based on BindingDB annotations or matching assay metadata, where an assay refers to a series of experiments testing different small molecules against the same protein target. The deduplicated records were then merged by UniProt ID. The final dataset contains approximately 2.5 million protein-ligand bioactivity records derived from about 67,000 distinct assays.

#### 3.3.2. Data Sampling

We developed a quality-prioritized sampling strategy to select training triplets from the bioactivity data. First, we ranked all assays by their data source reliability. Following this quality ranking, we proceeded to select assays in descending order of quality while simultaneously maximizing coverage of distinct UniProt IDs and ensuring no overlap with test set proteins. This process resulted in the selection of approximately 8000 high-quality assays. From each chosen assay we then extracted four distinct protein-ligand triplets, ultimately generating a balanced training set containing 30000 carefully curated data points that maintain both data quality and target diversity.

## 4. Experiments

### 4.1. Experiment Settings

#### Benchmark Datasets

We evaluate our methods on two benchmark datasets: **JACS 8** and **Merck FEP**. The JACS 8 dataset (Wang et al., 2015) consists of eight high-quality congeneric series extracted from real-world lead optimization projects. It is designed to benchmark fine-grained affinity ranking within structurally similar ligands. The Merck FEP benchmark (Schindler et al., 2020) contains ΔΔ*G* measurements across eight pharmaceutically relevant targets with greater structural diversity and noise, derived from highthroughput FEP benchmarking studies. For both datasets, we adopt the FEP+ setup, where each *edge* corresponds to a chemically meaningful ligand pair selected based on structural similarity, synthetic feasibility, and SAR significance. These pairs reflect realistic decisions encountered in lead optimization pipelines, where precise molecular ranking is more impactful.

#### Metrics

For each target, we evaluate model performance using Accuracy and Area Under the ROC Curve (AUC), based on the prediction of pairwise preferences between edges. Both metrics are computed per target and then averaged across all targets to obtain the final performance.

#### Baselines

Multiple baselines are compared, including physics-based methods FEP+(Wang et al., 2015) and MM-GB/SA(Lyne et al., 2006; Genheden & Ryde, 2015), which estimate binding affinities via simulation and energy approximations. Deep learning-based methods include PBCNet(Yu et al., 2023), EHIGN(Yang et al., 2024), and LigUnity(Feng et al., 2025). Finally, Protenix-ipTM uses AlphaFold3-style co-folding to extract interface confidence scores as affinity proxies(Shamir & London, 2025).

### 4.2. Experiment Results

Table 1 summarizes the performance of all models on the JACS and Merck datasets. On the JACS benchmark, our zero-shot method **AlphaRank**_**zero-shot**_ achieves a substantial improvement over the direct **Protenix-ipTM** baseline, boosting Accuracy from 0.5679 to 0.7048. This highlights that AlphaFold3-style confidence scores (e.g., ipTM) alone are insufficient for binding affinity ranking. Instead, our zero-shot approach introduces a biologically grounded inductive bias through competitive co-folding of ligand pairs with the same protein, effectively enabling relative ranking without additional supervision.

**Table 1.**
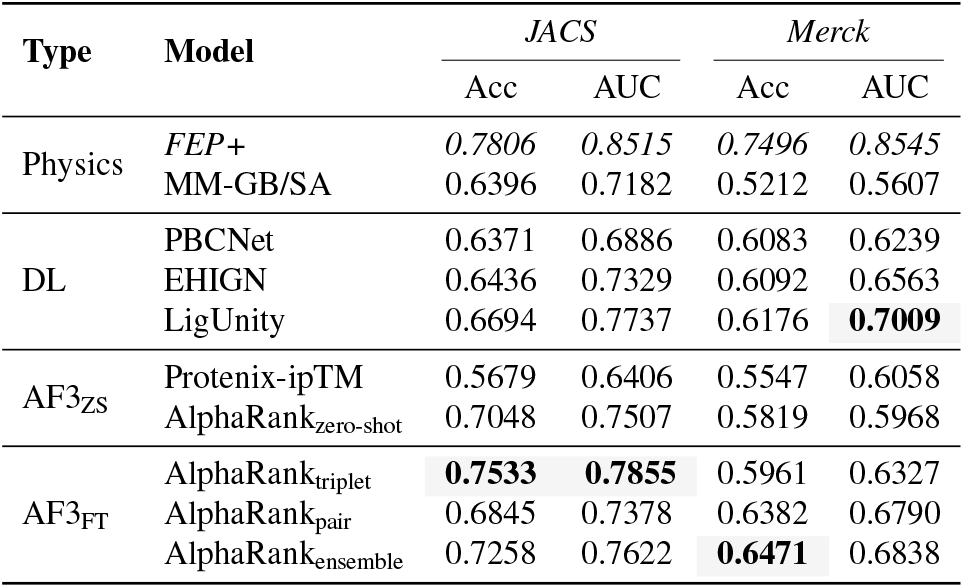
Performance comparison across different models. Models are grouped by type: physics-based methods, deep learning baselines, AF3 zero-shot, and finetuned AF3 features. **Bold** indicates the best learning-based performance on each dataset.

Furthermore, our fine-tuned models show consistent gains. The **AlphaRank**_**triplet**_ model achieves 0.7533 Accuracy and 0.7855 AUC on JACS, establishing a new state-of-the-art among learning-based methods. On Merck, the **AlphaRank**_**pair**_ model attains the highest Accuracy (0.6382) among individual learning-based approaches. When combined, the **AlphaRank**_**ensemble**_ model achieves the best overall performance, with 0.7258 Accuracy on JACS and 0.6471 on Merck, outperforming all deep learning baselines and approaching the accuracy of the physics-based gold standard **FEP+**. We believe the smaller margin on Merck attributes to its high-throughput screening origin, which introduces greater experimental noise and chemical diversity.

### 4.3. Analysis

Through our analysis, we found that two factors significantly impact the final ranking performance: the range of absolute ΔΔ*G* values and the average pLDDT scores during co-folding. As shown in Figure 2(a), for the Merck dataset, performance improves markedly as the magnitude of |ΔΔ*G*| increases. This suggests that the model is more reliable when ranking ligand pairs with larger affinity differences—an intuitive outcome, as such cases present clearer structural and energetic distinctions that are easier to capture. Interestingly, although our **AlphaRank**_**zero-shot**_ gives comparable AUC to the AlphaFold ipTM baseline on the full Merck set, it achieves noticeably higher AUC (0.7204 vs. 0.6766) when restricted to pairs with |ΔΔ*G*| *>* 1.0, indicating that its advantage becomes more pronounced in high-confidence ranking scenarios, particularly relevant for lead optimization, where correctly identifying chemical modifications that yield substantial affinity improvements is most critical. In contrast, performance on the JACS dataset remains relatively stable across different ΔΔ*G* intervals, suggesting that the model is well calibrated for subtle affinity changes, likely due to the congeneric nature and consistent SAR trends within JACS compound series.

In Figure 2(b), we observe a strong correlation between average pLDDT and model accuracy for Merck targets, while this correlation is notably weaker for JACS. This implies that pLDDT, a proxy for structural confidence, can serve as a useful indicator of model reliability on structurally diverse or noisier targets, such as those in Merck. As a result, pLDDT may be employed as a practical uncertainty signal to anticipate model performance on unseen targets, improving the interpretability and robustness of our approach.

**Figure 1.**
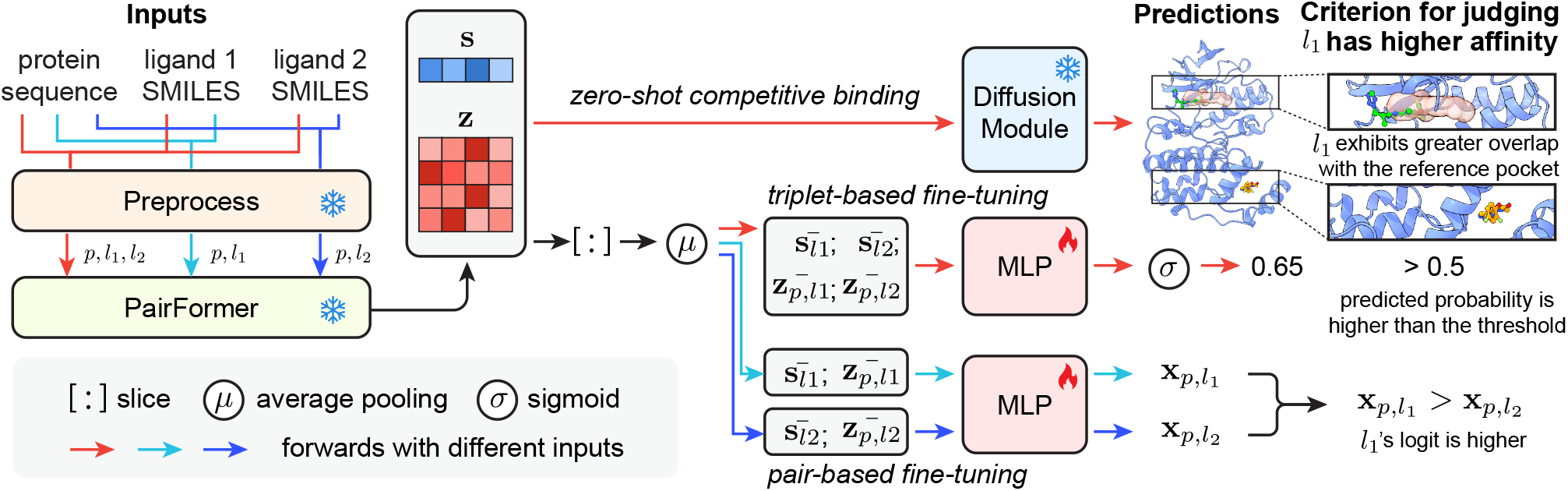
Illustration of AlphaRank with three affinity ranking strategies including zero-shot competitive binding, triplet-based fine-tuning, and pair-based fine-tuning.

**Figure 2.**
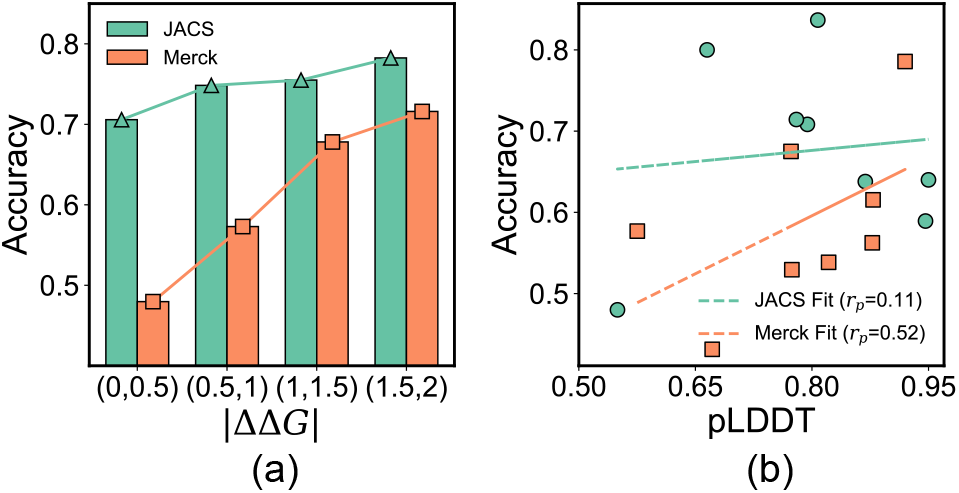
(a) Model performance grouped by the magnitude of affinity difference (|ΔΔ*G*|) label. (b) Relationship between model accuracy and average pLDDT scores across targets.

## 5. Conclusion and Future Work

In this paper, we present AlphaRank, the first framework to leverage AlphaFold3-style models for accurate affinity ranking by exploiting their capabilities in interaction modeling and co-folding prediction. The method incorporates a zero-shot setting based on competitive binding, and further enhance performance through fine-tuning on PairFormer-derived features. AlphaRank achieves state-of-the-art results on the JACS and Merck dataset, demonstrating the effectiveness of AlphaFold3-like model for lead optimization. Looking ahead, we plan to improve AlphaRank with smarter pair sampling strategies, integration of diffusion-based features, and end-to-end co-training. Our goal is to approach the accuracy of FEP+ while significantly reducing computational cost, making accurate affinity ranking more scalable and practical for drug discovery.

## Notes

### Competing Interest Statement

The authors have declared no competing interest.

## References

Abramson, J., Adler, J., Dunger, J., Evans, R., Green, T., Pritzel, A., Ronneberger, O., Willmore, L., Ballard, A. J., Bambrick, J., et al. Accurate structure prediction of biomolecular interactions with alphafold 3. Nature, 630(8016):493–500, 2024.

Burges, C., Shaked, T., Renshaw, E., Lazier, A., Deeds, M., Hamilton, N., and Hullender, G. Learning to rank using gradient descent. In Proceedings of the 22nd International Conference on Machine Learning - ICML ‘05, pp. 89–96, 2005.

Feng, B., Liu, Z., Yang, M., Zou, J., Cao, H., Li, Y., Zhang, L., and Wang, S. A foundation model for proteinligand affinity prediction through jointly optimizing virtual screening and hit-to-lead optimization. bioRxiv, pp. 2025–02, 2025.

Genheden, S. and Ryde, U. The mm/pbsa and mm/gbsa methods to estimate ligand-binding affinities. Expert opinion on drug discovery, 10(5):449–461, 2015.

Gilson, M. K., Liu, T., Baitaluk, M., Nicola, G., Hwang, L., and Chong, J. BindingDB in 2015: A public database for medicinal chemistry, computational chemistry and systems pharmacology. Nucleic Acids Research, 44(D1): D1045–D1053, January 2016. ISSN 0305-1048. doi: 10.1093/nar/gkv1072. URL https://doi.org/10.1093/nar/gkv1072.

Lyne, P. D., Lamb, M. L., and Saeh, J. C. Accurate prediction of the relative potencies of members of a series of kinase inhibitors using molecular docking and mm-gbsa scoring. Journal of Medicinal Chemistry, 49(16):4805–4808, 2006. doi: 10.1021/jm060522a. URL https://doi.org/10.1021/jm060522a. PMID: 16884290.

Mendez, D., Gaulton, A., Bento, A. P., Chambers, J., De Veij, M., Félix, E., Magariños, M. P., Mosquera, J. F., Mutowo, P., Nowotka, M., Gordillo-Marañoń, M., Hunter, F., Junco, L., Mugumbate, G., Rodriguez-Lopez, M., Atkinson, F., Bosc, N., Radoux, C. J., Segura-Cabrera, A., Hersey, A., and Leach, A. R. ChEMBL: towards direct deposition of bioassay data. Nucleic Acids Research, 47(D1):D930–D940, 2018. ISSN 0305-1048. doi: 10.1093/nar/gky1075.

Schindler, C. E., Baumann, H., Blum, A., Bose, D., Buchstaller, H.-P., Burgdorf, L., Cappel, D., Chekler, E., Czodrowski, P., Dorsch, D., et al. Large-scale assessment of binding free energy calculations in active drug discovery projects. Journal of Chemical Information and Modeling, 60(11):5457–5474, 2020.

Shamir, Y. and London, N. State-of-the-art covalent virtual screening with alphafold3. bioRxiv, pp. 2025–03, 2025.

Team, B. A. A., Chen, X., Zhang, Y., Lu, C., Ma, W., Guan, J., Gong, C., Yang, J., Zhang, H., Zhang, K., et al. Protenix-advancing structure prediction through a comprehensive alphafold3 reproduction. bioRxiv, pp. 2025–01, 2025.

Wang, L., Wu, Y., Deng, Y., Kim, B., Pierce, L., Krilov, G., Lupyan, D., Robinson, S., Dahlgren, M. K., Greenwood, J., et al. Accurate and reliable prediction of relative ligand binding potency in prospective drug discovery by way of a modern free-energy calculation protocol and force field. Journal of the American Chemical Society, 137(7): 2695–2703, 2015.

Yang, Z., Zhong, W., Lv, Q., Dong, T., Chen, G., and Chen, C. Y.-C. Interaction-based inductive bias in graph neural networks: enhancing protein-ligand binding affinity predictions from 3d structures. IEEE Transactions on Pattern Analysis and Machine Intelligence, 2024.

Yu, J., Li, Z., Chen, G., Kong, X., Hu, J., Wang, D., Cao, D., Li, Y., Huo, R., Wang, G., et al. Computing the relative binding affinity of ligands based on a pairwise binding comparison network. Nature Computational Science, 3(10):860–872, 2023.

